# Interleukin-6 is critical in the development of *gcn2-*mutation associated pulmonary vascular disease in mice

**DOI:** 10.1101/2025.04.14.648729

**Authors:** Max Schwiening, Qingyue Gao, Mark Southwood, Alexi Crosby, Stephen Moore, Jose Antonio Valer, Niki Veale, Benjamin J. Dunmore, Paul D. Upton, Roger Thompson, Nicholas W. Morrell, Stefan J. Marciniak, Elaine Soon

## Abstract

Biallelic mutations in eukaryotic translation initiation factor 2 α kinase 4, *EIF2AK4* (which encodes general control nonderepressible 2, GCN2) underpin heritable forms of pulmonary veno-occlusive disease (PVOD), a rare and fatal form of pulmonary hypertension. The mechanisms linking these are mostly uncharacterised. We demonstrate for the first time that homozygous loss of *gcn2* is sufficient to cause mild pulmonary hypertension in mice. Single-cell transcriptomics of mouse lungs identified adventitial fibroblasts as having the greatest GCN2-dependent transcriptional differences, implicating them as key players in this model of PVOD. The most significantly upregulated pathways in *gcn2^-/-^* adventitial fibroblasts were inflammatory. Therefore, we went on to demonstrate a pro-inflammatory phenotype in *gcn2^-/-^* mouse embryonic fibroblasts and *gcn2^-/-^* mice. In a novel murine model of pulmonary hypertension induced by exposure to mitomycin C, deletion of interleukin-6 rescued the pulmonary vascular phenotype. When chronically exposed to lipopolysaccharide, the pulmonary hypertensive phenotype of *gcn2^-/-^* mice is exaggerated. Genetic ablation of interleukin-6 completely rescues both the baseline and LPS-exaggerated pulmonary hypertensive phenotype. Targeting *Il6*-dependent pathways may be useful in treating this deadly disease.

## Introduction

Pulmonary arterial hypertension (PAH) is an umbrella term describing diseases characterised by an increase in mean pulmonary artery pressures, with a normal left heart[1]. Pulmonary veno-occlusive disease (PVOD) is a rare but deadly form of pulmonary arterial hypertension. The histological hallmark of PVOD is the progressive obliteration of small pulmonary veins and venules by fibrous intimal thickening. The clinical presentation is one of worsening breathlessness on exertion as the increasing pulmonary vascular resistance results in right heart failure and death. Comparatively little is known about the pathophysiology of PVOD and these patients are generally treated empirically with medications licensed for PAH, with particularly poor outcomes. Most patients with PVOD die within 2-3 years of diagnosis without lung transplantation[2]. PVOD has been reported following exposure to various environmental toxins and drugs, such as mitomycin C[3, 4], organic solvents[5]; and autoimmune/ inflammatory conditions[6, 7].

Biallelic mutations in eukaryotic translation initiation factor 2 α kinase 4 or *EIF2AK4* (which encodes general control nonderepressible 2, GCN2), were identified as causative in familial forms of PVOD in 2014[8]. Mutations were found in all familial cases and 25% of sporadic PVOD. These patients were also younger at diagnosis and had a worse prognosis than patients with idiopathic pulmonary hypertension[9]. Levels of GCN2 were also shown to be reduced in sporadic PVOD as well as in heritable and idiopathic PAH[10]. GCN2 is a serine/threonine protein kinase, which phosphorylates the α-subunit of the translation initiation factor eIF2, activating the Integrated Stress Response[11]. This results in the reduction of translation initiation in general and the translation of specific messenger RNAs to either restore homeostasis, or, if overwhelming, to trigger apoptosis. GCN2 responds specifically to amino acid deprivation, and canonically it has been regarded as a key regulator of metabolic stress.

There is now a robust body of evidence linking GCN2 to control of inflammation and immune processes in the gut [12, 13] and the central nervous system[14]. *Gcn2-*deficient mice showed an exaggerated inflammatory response upon exposure to 2% dextran sodium sulphate, which triggers an inflammatory colitis. This was characterized by Ravindran *et al*[12], who proved that *gcn2^-/-^* mice showed greater weight loss, higher levels of IL-17, more severe intestinal histological changes and increased intestinal permeability post-DSS exposure. Similarly, *gcn2-*deficient mice developed greater weight loss, higher levels of TNF-α and pancreatitis-associated protein mRNA expression, more severe acinar cell histological changes post-exposure to asparaginase (a chemotherapy agent commonly used in acute lymphoblastic leukaemia)[13].

This effect is not limited to the gut, as GCN2 has been shown to play a significant role in controlling neuroinflammation [14, 15] and the haematopoietic/immune cell population [16, 17]. The pattern is slightly different in the central nervous system, with *gcn2^-/-^* mice showing a failure to resolve neuroinflammation, specifically a model of experimental autoimmune encephalomyelitis induced by immunization with myelin oligodendrocyte glycoprotein peptide 33-55[14], which is a model for multiple sclerosis. The *gcn2-*deficient mice presented with a persistent paralysis, while the majority of similarly immunized wild-type littermates recovered. This susceptibility was transferrable with *gcn2^-/-^* bone marrow and was replicated with T-cell specific GCN2 deletion.

Pulmonary arterial hypertension in general also has a strong association with inflammatory phenotypes in cell lines, animal models and in patients. Idiopathic and heritable PAH patients not only have higher levels of pro-inflammatory cytokines, but these levels also predict mortality [18, 19]. Chronic exposure to pro-inflammatory insults replicates the pulmonary arterial hypertension phenotype in mice bearing mutations in bone morphogenetic protein receptor type 2, *bmpr2* [20]; the other main genetic cause of pulmonary arterial hypertension. Transgenic rodents overexpressing interleukin-6 are more susceptible to hypoxia-driven pulmonary vasoconstriction, while mice with impairments in IL-6 signalling are protected from this [21-23]. Therefore, we hypothesised that *gcn2* mutations lead to a pro-inflammatory state in the lung, which triggers and propagates the pathogenesis of pulmonary vascular disease; and that interruption of the pro-inflammatory pathways involved would abrogate the pulmonary vascular response.

## Results

### *Gcn2* deficiency in mice produces a mild pulmonary hypertensive phenotype

Wild-type, *gcn2^+/-^* and *gcn2^-/-^* mice aged between 4.5 and 6 months were subjected to right and left heart catheterization and echocardiography to characterise their cardiovascular system at baseline. The *gcn2^+/-^* and *gcn2^-/-^* mice both displayed a mild increase in their right ventricular systolic pressures (RVSP) compared to wild-type littermates (mean RVSP of 24.3±3.5 mmHg in wild-type, *versus* 27.4±4.1 mmHg in *gcn2^+/-^* and 28.2±4.1mmHg in *gcn2^-/-^* mice, Fig.1A). This was associated with a concomitant increase in the right ventricular mass indexed to body weight (RV/BW, Fig.1B). The Fulton index (right ventricular mass indexed to the mass of the left ventricle and septum, RV/LV+S) did not differ between wild-type, *gcn2^+/-^* and *gcn2^-/-^* mice (Fig.1C). The left ventricular systolic pressures (LVSP, Fig.1D) also did not differ between the genotypes. However, there was an increase in the left ventricular mass indexed to body weight (LV/BW, Fig.1E) in both *gcn2^+/-^* and *gcn2^-/-^* mice. The body weight of these mice was not different between genotypes (Fig.1F). The cardiac index (defined as cardiac output/body surface area), and the left ventricular anterior wall thickness in diastole (LVAW:D) as measured by echocardiography, did not differ between genotypes (Fig.1G and Fig.1H). There was a non-significant increase in the left ventricular posterior wall in diastole (LVPW:D, Fig.1I) in the *gcn2^+/-^*and *gcn2^-/-^* mice. When the small pulmonary vessels (50-200µm in diameter) of these mice were examined, there was a change in the pattern of vessel muscularisation, as shown by the degree of α-smooth muscle actin staining (Fig.1J). Wild-type mice had predominantly unmuscularised vessels, while both *gcn2^+/-^* and *gcn2^-/-^* mice had a shift towards increasing muscularisation with predominantly partially muscularised vessels (Fig.1K).

**Figure 1:**
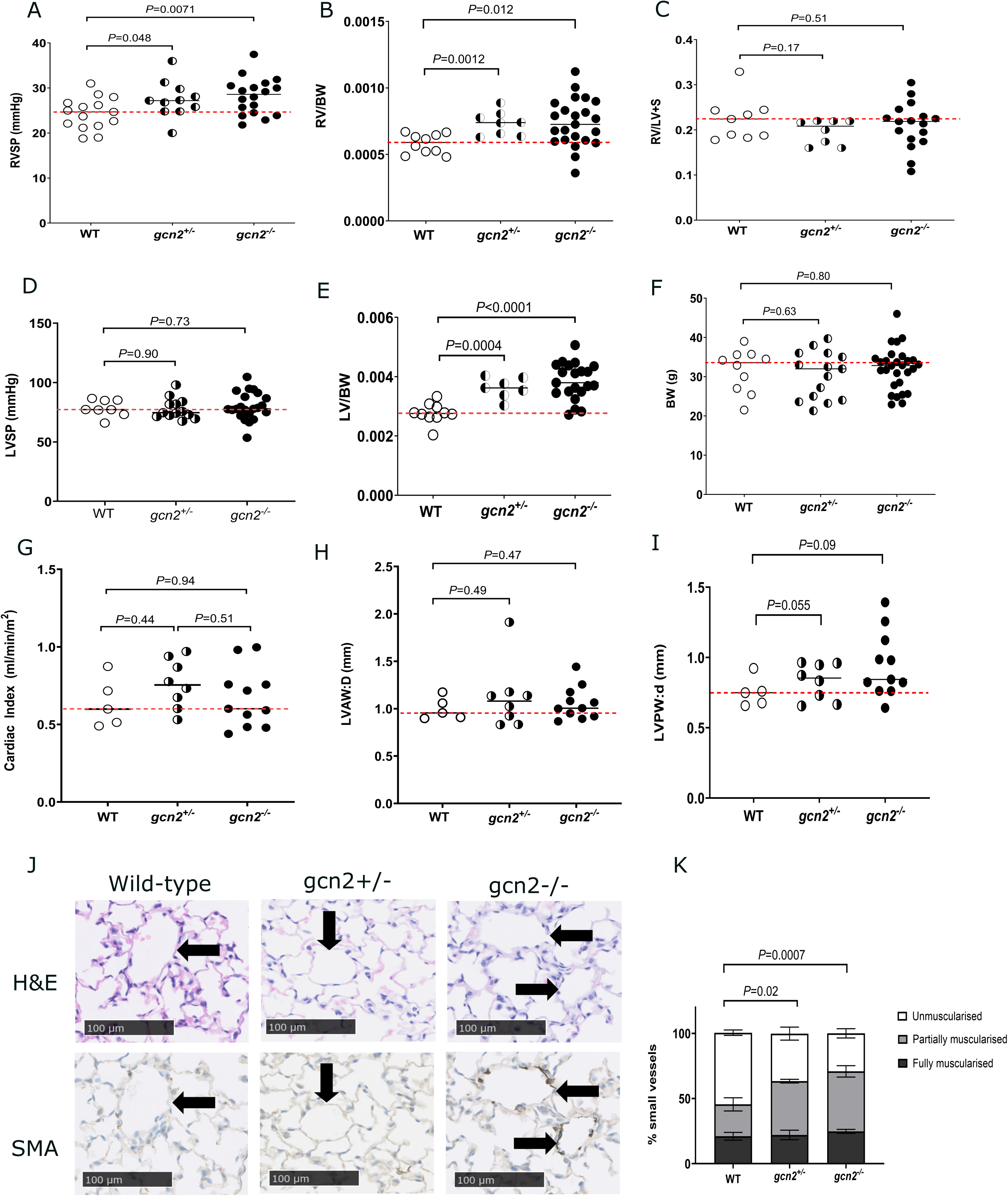
Characterisation of the *gcn2*-deficient mouse at baseline. Panels A-C show the right ventricular systolic pressure (RVSP, panel A), the right ventricular mass indexed to body weight (RV/BW, panel B) and the right ventricular mass indexed to septal and left ventricular mass (RV/LV+S, panel C) in mice which are wild-type, heterozygous and homozygous for a deletion in *gcn2* (wild-type, *gcn2^+/-^* and *gcn2^-/-^)*. Panels D-F show the left ventricular systolic pressure (LVSP, panel D), left ventricular mass indexed to body weight (LV/BW, panel E) and the body weight (panel F) of wild-type, *gcn2^+/-^* and *gcn2^-/-^* mice. For panels A and D, the right and left ventricular pressures have been measured by right and left cardiac catheterisation. Panels G-I show the cardiac index (defined as cardiac output per body surface area, panel G), the left ventricle anterior wall thickness measured at diastole (LVAW:D, panel H) and the left ventricle posterior wall thickness measured at diastole (LVPW:D, panel I) of wild-type, *gcn2^+/-^* and *gcn2^-/-^* mice measured by echocardiography. Panel J shows representative histological sections of the lungs of wild-type, *gcn2^+/-^* and *gcn2^-/-^* mice, stained with haematoxylin and eosin (H&E) and smooth muscle actin (SMA). Panel K shows the quantification of non-muscularised, partially muscularised and fully muscularised vessels. In panels (A-I) the median is shown, and comparisons have been made using unpaired t-tests (for parametric data) or Mann-Whitney tests (for non-parametric data). For panel K comparisons of the distribution of non-muscularised: partially muscularised: fully muscularised vessels in wild-type, *gcn2^+/-^* and *gcn2^-/-^* mouse lungs were made using Fisher’s contingency test.

### Single-cell RNA sequencing provides insights into the pathways and cell types responsible for the phenotype associated with *gcn2* deficiency

Our next step was to investigate which pathways might be responsible for the pulmonary vascular phenotype observed above. To do this in an unbiased manner, the lungs of *gcn2^-/-^* and wild-type littermates raised in the same cage were extracted and processed to generate single-cell suspensions for single cell RNA sequencing (scRNAseq, schema shown in Fig.2A). Isolation of some cell types, particularly immune cells, is more efficient than others and can skew datasets[24]. We therefore chose to sort cells by CD45 expression to generate CD45-positive populations (immune cells) and CD45-negative (all other cell types). The Uniform Manifold Approximation and Projection (UMAP) protocol was used for dimensionality reduction and cell clusters from the CD45-negative group (Fig.2B-C) and CD45-positive group (Supplementary Fig.1A-B) were identified by known marker genes.

**Figure 2:**
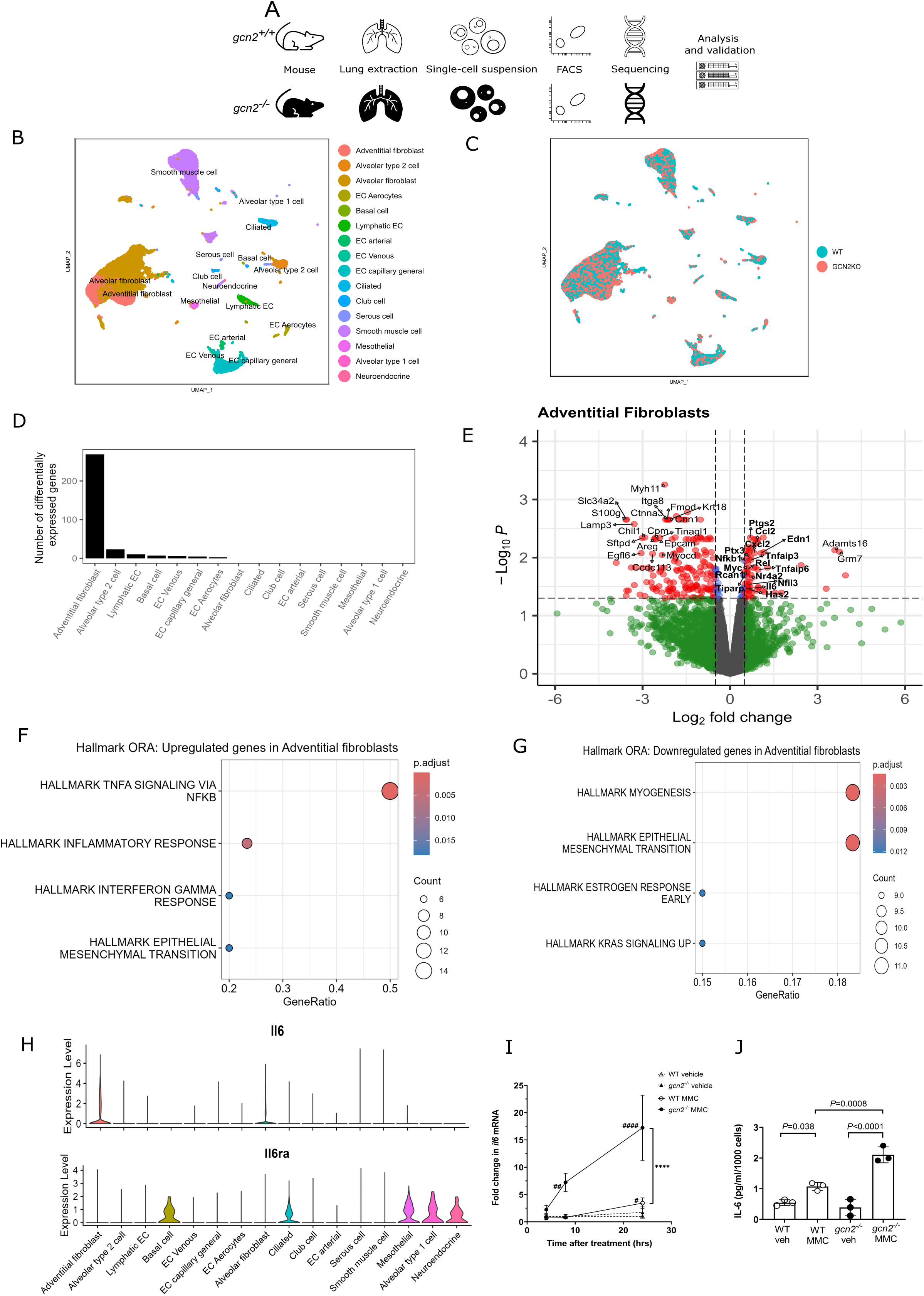
Single-cell RNA sequencing of wild-type and *gcn2-*deficient mouse lungs. Panel A shows a schematic for this experiment, in which pairs of age- and sex-matched littermates who were wild-type and *gcn2^-/-^* respectively were humanely killed, lungs collected and dissociated into single-cell suspensions, sorted for size, viability and CD45-expression by flow cytometry and sent for single cell RNA sequencing. Panels B and C show the Uniform Manifold Approximation and Projection graphs (UMAPs) of CD45-negative lung cells, with panel B depicting cell clusters and panel C showing contributions from the wild-type and *gcn2^-/-^* mice. Panel D shows the cell types from the CD45-negative group arranged in order by the number of genes differentially expressed between wild-type and *gcn2^-/-^* lung cells. Panel E shows a volcano plot of differentially expressed genes between genotypes in adventitial fibroblasts, with genes from upregulated inflammatory pathways identified by ORA in bold. Panels F and G shows pathways that were activated or suppressed in the *gcn2^-/-^* adventitial fibroblasts with respect to their wild-type counterparts. Panel H shows violin plots showing the relative expression of *Il6* and *Il6ra* in cell types from the CD45-negative dataset. The matching plots for the CD45-positive dataset is shown in Supplementary Figure 1, panels A-D. Panel I shows the levels of mRNA encoding IL-6 from wild-type and *gcn2^-/-^*MEFS after exposure to either media alone (DMEM+10%FBS) or 1µg/ml of MMC for 4, 8 and 24 hours. Panel J shows the quantification of IL-6 secreted into media from wild-type and *gcn2^-/-^* MEFS after exposure to either media (DMEM+10%FBS) or 1µg/ml of MMC for 24 hours. For panel H, a 2-way ANOVA was performed with Sidak’s multiple comparison testing. For panel I, a one-way ANOVA was performed with Sidak’s multiple comparison testing.

A pseudo-bulk quasi-likelihood model[25] was used to identify genes differentially expressed between wild-type and *gcn2^-/-^* cells for both the CD45-negative (Fig.2D) and CD45-positive (Supplementary Fig.1C) populations. Adventitial fibroblasts proved to be the cell type with the highest number of differentially expressed genes by a considerable margin, with 269 differently expressed genes compared to less than 25 for type 2 alveolar cells and lymphatic endothelial cells, which were the second and third ranked cell types. As adventitial fibroblasts appear to be a critical cell type, we examined the differentially expressed genes in this cluster in greater detail (volcano plot shown in Fig.2E). Over-representation analysis of differentially expressed genes reveals a highly significant upregulation of pathways associated with inflammatory signalling (Fig.2F-G). The inflammatory cytokine that was detected in this setting is *Il6.* This is intriguing as IL-6 has been closely associated with pulmonary arterial hypertension in patients[18, 19, 26] and rodent models[21-23].

We then interrogated the *Il6* gene expression in all cell types and found that adventitial fibroblasts demonstrated the greatest expression of *Il6* (Fig.2H and Supplementary Fig.1D). Expression of the cognate receptor *il6ra* was more diffuse with expression in basal cells, ciliated cells, mesothelial, alveolar type 1 cells, and neuroendocrine cells in the CD45-negative population and macrophages, dendritic cells, and neutrophils in the CD45-positive population (Supplementary Fig.1D). A table showing specific genes and the roles of their encoded proteins is shown in Supplementary Fig.1E.

To validate these results, we tested the hypothesis that *gcn2-*deficient fibroblasts would have a pro-inflammatory cytokine response by exposing wild-type and *gcn2^-/-^* mouse embryonic fibroblasts (MEFs) to mitomycin C (MMC) *in vitro* for 4, 8, and 24 hours (Fig.2I). Mitomycin C is a chemotherapeutic agent used in certain cancers and was specifically chosen as the trigger as it has the idiosyncratic side effect of causing pulmonary veno-occlusive disease which is clinically very similar to the idiopathic and *gcn2-*mutation related forms of PVOD[27]. We found that *Il6* mRNA was significantly elevated in both wild-type and *gcn2-*deficient MEFs treated with MMC at 24 hours compared to those in media alone. *Gcn2^-/-^* MMC-treated MEFs showed greater levels of *il6* mRNA at the 8 and 24-hour timepoints compared to wild-type MMC-treated MEFs. This was confirmed using ELISAs to measure IL-6 levels in the cultured media from these cells at the 24-hour time point, demonstrating that MMC treatment leads to synthesis and secretion of this cytokine; and that levels of secreted IL-6 are significantly increased in *gcn2-*deficient MEFs (Fig.2J).

### Genetic ablation of *il6* is protective in a murine model of mitomycin C induced pulmonary hypertension

To investigate whether *Il6* is pivotal in PVOD, we developed a mouse model of disease utilising mitomycin C (MMC) as the trigger. Wild-type or *il6^-/-^* mice aged 8-11 weeks were injected with cumulative doses of mitomycin-C totalling either 0.5mg/kg, 1.0 mg/kg, or 2.0mg/kg; or vehicle alone, and subjected to right heart catheterisation 3 weeks later (schematic shown in Fig.3A). Wild-type mice exposed to mitomycin-C developed an increase in their right ventricular mass indexed to left ventricular mass (Fulton index, Fig.3B), right ventricular systolic pressures (Fig.3C) and right ventricular mass indexed to body weight (Fig.3D). Strikingly, interleukin-6-deficient mice were protected from the development of elevated RVSP at all doses (Fig.3C). There was no change in the left ventricular mass indexed to body weight (Fig.3E) for both the wild-type and *Il6^-/-^* mice. Examination of lung sections revealed increased muscularisation of the smaller vessels in the wild-type mouse lungs chronically exposed to MMC (Fig.3F-G). These changes did not occur in *Il6^-/-^* mice.

**Figure 3:**
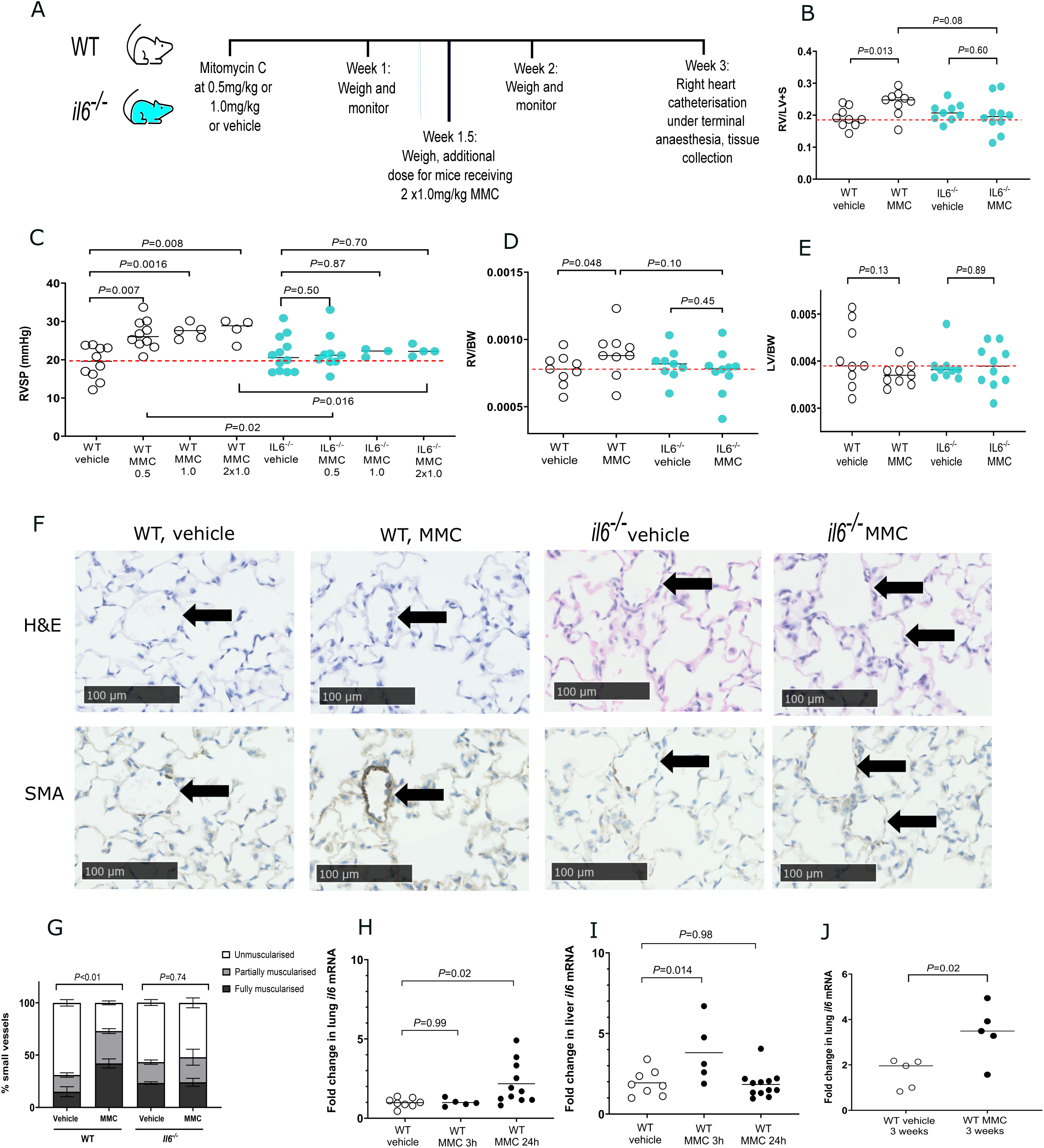
The characterisation of a murine model of mitomycin-C (MMC) induced PVOD. Panel A shows a schematic for this experiment, in which wild-type and *il6^-/-^* mice were exposed to either vehicle or 0.5mg/kg, 1.0mg/kg or 2.0mg/kg (in total) of MMC. Panel B shows the right ventricular mass indexed to left ventricular and septal mass (RV/LV+S). Panel C shows the right ventricular systolic pressure (RVSP) for all mice in the study. Panels (D-E) show the right ventricular mass indexed to body weight (RV/BW, panel D), and the left ventricular mass indexed to body weight (LV/BW, panel E) for wild-type and *il6^-/-^* mice exposed to vehicle or 0.5mg/kg of MMC. Panel F shows representative histological lung sections of wild-type and *il6^-/-^* mice from both the vehicle and MMC-treated arms, stained with haematoxylin and eosin (H&E) and smooth muscle actin (SMA). Panel G shows the quantification of non-muscularised, partially muscularised and fully muscularised vessels of vessels which are 50-200microns in diameter in a 10mm^2^ lung area section per mouse. Panels H and I show the levels of mRNA encoding IL-6 in wild-type mice after exposure to either vehicle or MMC for either 3 or 24hours in RNA, extracted from lung (panel H) and liver (panel I). Panel J shows the levels of lung mRNA encoding IL-6 in wild-type mice after exposure to either vehicle or MMC for 3 weeks. In panels (B, C, D, E, H, I, and J) the median is shown, and comparisons have been made using unpaired t-tests (for parametric data) or Mann-Whitney tests (for non-parametric data). In panel G the distribution of non-muscularised: partially muscularised: fully muscularised vessels has been compared between groups using Fisher’s contingency testing.

To determine if acute exposure to MMC elicited a specific increase in pulmonary IL-6, 6- to 8-week-old wild-type mice were exposed to 1.0mg/kg of mitomycin C or vehicle. They were euthanised 3- or 24-hours post-treatment and lung and liver tissue was collected. The mRNA encoding IL-6 in the lungs was increased at 24 hours by MMC treatment (Fig.3H). There was also an increase in liver IL-6 mRNA but this occurred earlier, at the 3-hour timepoint (Fig.3I). When wild-type mice were exposed to a single dose of mitomycin C and euthanised 3 weeks later, the increase in lung IL-6 mRNA is maintained (Fig.3J).

*Gcn2* deficiency is associated with a more marked inflammatory response to lipopolysaccharide. Chronic exposure to lipopolysaccharide induces a worse pulmonary hypertensive phenotype in *gcn2*-deficient mice.

We next sought to test if GCN2-deficiency would be associated with a general pro-inflammatory phenotype. Plasma samples were obtained from the National Cohort Study of Idiopathic and Heritable Pulmonary Arterial Hypertension. Our PVOD patients with biallelic mutations in *GCN2* had higher levels of plasma IL-6 compared to age- and sex-matched healthy volunteers (Fig.4A). Aged *gcn2^-/-^* mice displayed an increase in lung mRNA encoding IL-6 compared to their wild-type littermates (Fig.4B). This difference was further exaggerated after an intraperitoneal injection of 0.05mg/kg of lipopolysaccharide (LPS) for three hours (Fig.4C). Levels of keratinocyte-derived cytokine (KC, a mouse analogue of interleukin-8 that functions as a neutrophil chemoattractant) were elevated in a similar manner (Fig.4D). We hypothesised that since an acute exposure to LPS exaggerated difference in pro-inflammatory cytokines between wild-type and *gcn2^-/-^* mice, chronic exposure could exaggerate differences in the cardio-pulmonary phenotype. To test this, 8–10-week-old wild-type and *gcn2^-/-^* mice were injected with either vehicle or LPS thrice weekly for six weeks (schematic shown in Fig.4E). *Gcn2^-/-^* mice demonstrated a further increase in their right ventricular systolic pressures after this treatment (Fig.4F). Neither right ventricular mass indexed to body mass (RV/BW) nor left ventricular mass indexed to body mass (LV/BW) changed appreciably during this experiment in either genotype (Figs.4G-2H).

**Figure 4:**
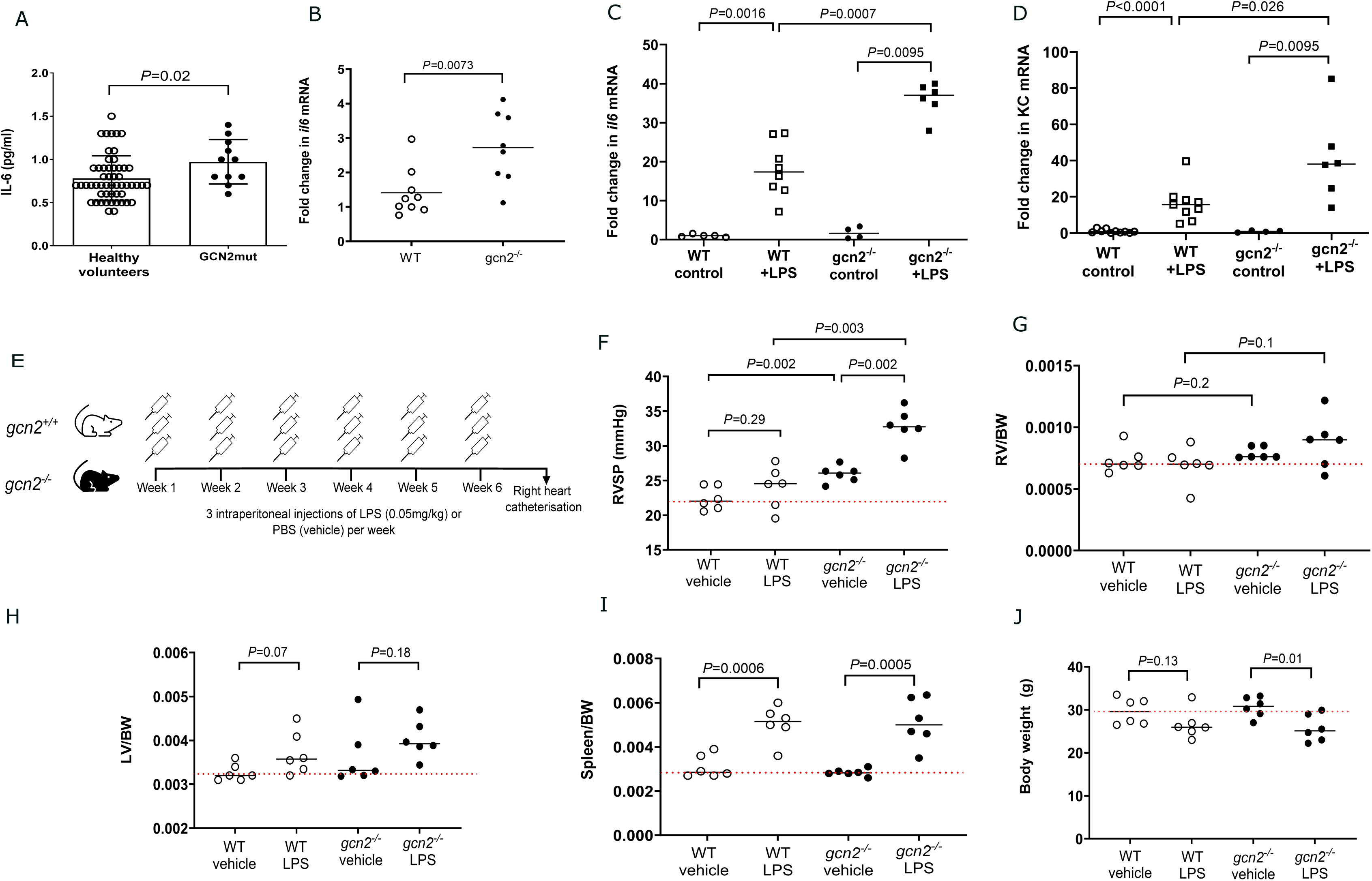
*Gcn2*-deficiency is associated with greater levels of pro-inflammatory cytokines at baseline and after stimulation with lipopolysaccharide, LPS. Panel A shows the levels of plasma IL-6 from healthy volunteers and PVOD patients bearing biallelic GCN2 mutations. Panel B shows the levels of mRNA encoding IL-6 from the lungs of baseline aged wild-type and *gcn2^-/-^* mice. Panels C and D show the fold change in lung mRNA encoding IL-6 (panel C) and KC (panel D) after exposure to either vehicle or 0.05mg/kg of LPS in young wild-type and *gcn2^-/-^* mice. KC is a mouse analogue of interleukin-8. Panel E shows a schematic for the chronic LPS experiment, in which wild-type, *gcn2^+/-^* and *gcn2^-/-^* mice were exposed to either vehicle or LPS thrice weekly for 6 weeks. Panels F-J show the results from chronic exposure to either vehicle or LPS. Panel F shows the right ventricular systolic pressures (RVSP), panel G the right ventricular mass indexed to body weight (RV/BW) and panel H the left ventricular mass indexed to body weight (LV/BW) of these mice. Panel I shows the spleen indexed to body weight (spleen/BW) and panel J the body weights of these mice. In data panels the median is shown, and comparisons have been made using unpaired t-tests (for parametric data) or Mann-Whitney tests (for non-parametric data). For panels B, C, and D, a wild-type mouse either at baseline or exposed to vehicle has been selected as the baseline control.

As expected, all genotypes displayed an LPS-induced increase in spleen weight indexed to body weight (spleen/BW, Fig.4I) compared to vehicle-treated mice of the same genotype, proving that a systemic inflammatory response had been elicited by the chosen regimen of LPS. The wild-type mice receiving 6 weeks of intraperitoneal LPS were not significantly lighter than wild-type mice receiving vehicle alone, while the *gcn2^-/-^* mice receiving LPS weighed significantly less than *gcn2^-/-^* treated with vehicle alone (Fig.4J).

### Genetic ablation of *il6* abrogates the pulmonary hypertensive phenotype both at baseline and after exposure to lipopolysaccharide

To test the role of IL-6 in development of the pulmonary hypertensive phenotype in baseline *gcn2-*deficient mice, we generated mice deficient in both *gcn2* and *Il6*. *II6^-/-^* and *gcn2^-/-^Il6^-/-^* mice were aged to 4.5-6 months and then subjected to right heart catheterisation (Supplementary Figure 2). Genetic loss of *Il6* effectively reverses the increased RVSP associated with *gcn2-*loss at baseline (Supplementary Fig.2A). The *Il6^-/-^* mice did not have significantly different RVSPs compared with wild-type animals at baseline. Interestingly, both *Il6^-/-^* and *gcn2^-/-^Il6^-/-^* mice had increased right ventricular mass indexed to body weight (RV/BW, Supplementary Fig.2B) and left ventricular mass indexed to body weight (LV/BW, Supplementary Fig.2C) compared to the wild-type at baseline. However, neither the RV/BW nor the LV/BW of these mice were different from the *gcn2^-/-^* mouse. When body weights were compared, neither the *Il6^-/-^* nor the *gcn2^-/-^il6^-/-^* mouse were significantly lighter than the wild-type (Supplementary Fig.2D). We note that *gcn2^-/-^Il6^-/-^* mice were significantly lighter than the *gcn2^-/-^* mice. Histological examination revealed that the lungs of *Il6^-/-^* and *gcn2^-/-^Il6^-/-^* mice resembled those of the wild-type, with predominantly unmuscularised small pulmonary vessels (Supplementary Fig.2E-F).

To test if IL-6 continues to be significant in LPS-induced pulmonary vascular disease, we injected 8–to-11-week-old *gcn2^-/-^Il6^-/-^* animals, and wild-type, *gcn2^-/-^*, and *Il6^-/-^* controls, with either LPS or vehicle thrice weekly for 6 weeks (schematic shown in Fig.5A). Genetic ablation of *il6* abrogated the right ventricular systolic pressure increase in both vehicle and LPS-exposed mice (Fig.5B). Neither right ventricular mass indexed to body mass (RV/BW) nor left ventricular mass indexed to body mass (LV/BW) changed appreciably during the experiment in most genotypes (Figs.5C-D). Immunohistochemical examination of lungs of *gcn2^-/-^* mice revealed increased muscularisation of the smaller lung vessels (50-200μm) as shown by α-smooth muscle actin (SMA) staining in the *gcn2^-/-^* mice chronically exposed to LPS. These changes did not occur in the wild-type, *gcn2^-/-^il6^-/-^* or *il6^-/-^* mice (Fig.5E-F).

**Figure 5:**
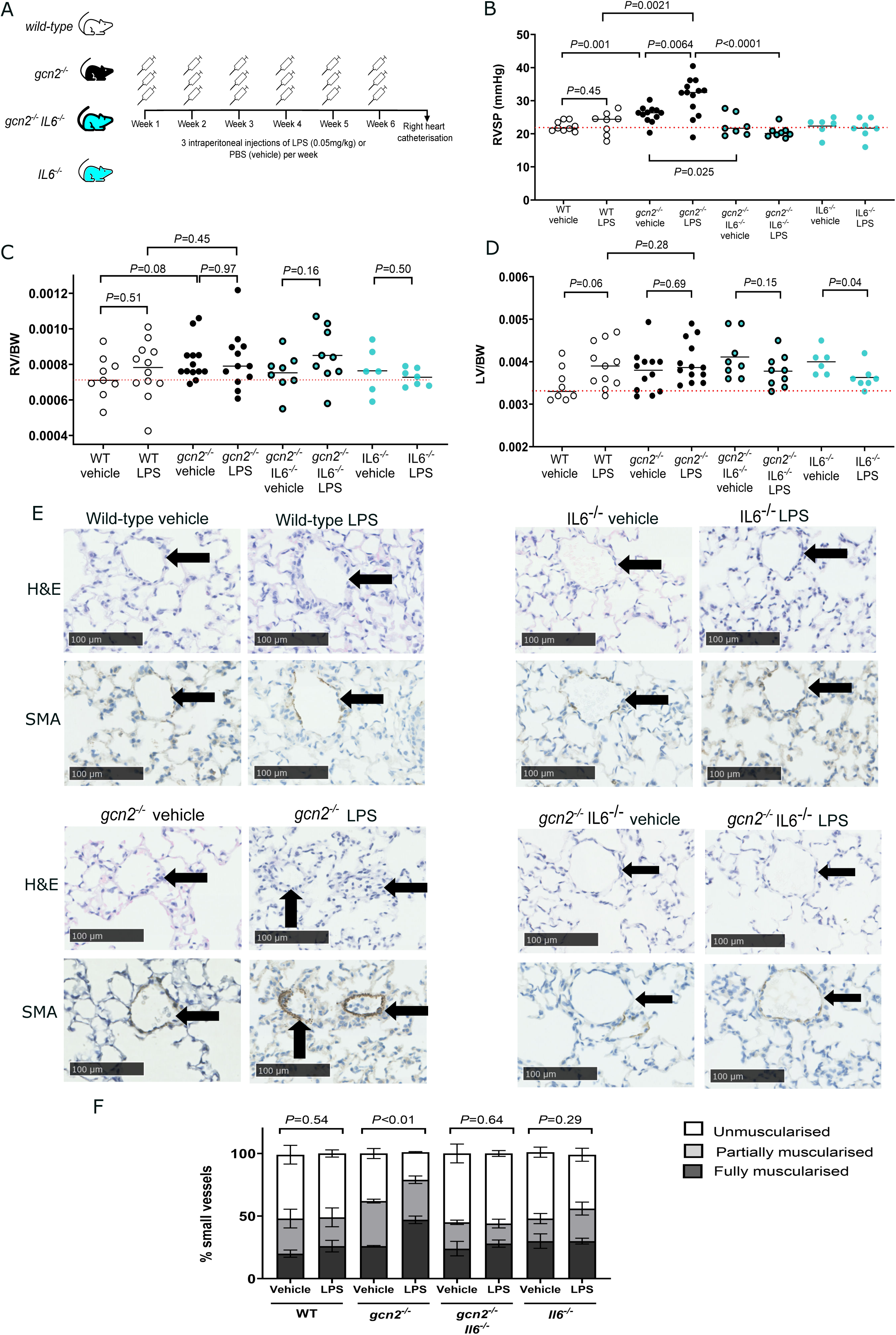
The baseline and exaggerated RVSP response to LPS in *gcn2-*deficient mice is abrogated by genetic loss of IL-6. Panel A shows a schematic for this experiment, in which wild-type, *gcn2^-/-^*, *Il6^-/-^* and *gcn2^-/-^ Il6^-/-^* mice were exposed to either vehicle or 0.05mg/kg of LPS thrice weekly for 6 weeks. Panels B-D show the right ventricular systolic pressure (RVSP, panel B), the right ventricular mass indexed to body weight (RV/BW, panel C) and the left ventricular mass indexed to body weight (LV/BW, panel D). Panels E-F show representative histological sections of the lungs of wild-type, *gcn2^-/-^*, *Il6^-/-^* and *gcn2^-/-^ Il6^-/-^* mice from both the vehicle and LPS-treated arms, stained with haematoxylin and eosin (H&E) and smooth muscle actin (SMA, panel E) and the quantification of non-muscularised, partially muscularised and fully muscularised vessels (panel F). In panels (B-D) the median is shown, and comparisons have been made using unpaired t-tests (for parametric data) or Mann-Whitney tests (for non-parametric data). In panel F the distribution of non-muscularised: partially muscularised: fully muscularised vessels has been compared between groups using Fisher’s contingency testing.

## Discussion

We have demonstrated for the first time that interleukin-6 and its pathways are critical for the development of pulmonary veno-occlusive-like disease in mice. To do this we developed two new mouse models of pulmonary veno-occlusive disease where none previously existed. The first model is a murine model with homozygous *gcn2* deletions that is analogous to *gcn2-*mutation bearing PVOD patients, who are either homozygotes or compound heterozygotes. The second model utilizes a chemotherapy agent, mitomycin C, which also directly reflects disease observed in patients. Our current theory is that *gcn2-*deficiency confers an innate susceptibility to inflammatory insults and that the resultant interleukin-6 driven inflammatory cascade leads to pulmonary vascular remodelling and eventually pulmonary vascular disease (Figure 6).

**Figure 6:** Graphical summary of hypothesis of PVOD development. Statements in red are evidenced in this manuscript, data in black are from the following references, and statements in blue are true of IL-6 in general but are not directly linked to GCN2 deficiency.

There is increasing evidence that this theory is both plausible and consistent. IL-6 is implicated in multiple rodent models of pulmonary hypertension including hypoxia-induced, monocrotaline-induced and Sugen-5416 and hypoxia models [21-23]. Independent of PAH models, IL-6 also mediates adverse cardiac remodelling[28] and a vasoconstrictive profile [29] in rodents. We have shown for the first time that IL-6 is elevated in the serum of GCN2-associated PVOD patients. IL-6 levels are an independent risk factor for adverse outcomes in heart failure patients with preserved ejection failure [30, 31]. Circulating IL-6 levels have been significantly positively correlated with both pulmonary arterial hypertension and systolic blood pressure elevation [32, 33].

Loss of GCN2 has been shown in rodents to lead to disruption of T-cell development, promotion of pro-inflammatory cytokine production and to impede recovery from immune and inflammatory insults [12, 34-36]. In rats, *gcn2* deficiency was strongly associated with a pulmonary inflammatory response, triggered specifically by acute asparagine and glutamine deprivation[36]. This was proven by expression of pro-inflammatory genesets, increased plasma cytokine levels, and infiltration of the lung perivascular space by inflammatory cells. More specifically, upregulation of interleukin-6 was observed in *gcn2-*deficient rats exposed to asparaginase at both the messenger RNA level (transcriptome analysis and lung quantitative PCR), and at the protein level (plasma levels). The investigators were unable to test the hypothesis that this would lead to pulmonary vascular disease since the *gcn2-*deficient rats were so unwell that they had to be euthanised 3 days post-exposure, which is insufficient time to allow for pulmonary vascular remodelling. This is the reason we have deliberately chosen to test lower doses of mitomycin-c and LPS, as we were not seeking to explore acute toxic or septic models but rather to test the premise that low level inflammation leads to pulmonary vascular remodelling in the context of GCN2 deficiency.

The literature also supports the premise that *gcn2-*deficiency leads to a more severe inflammatory phenotype in the context of experimental inflammatory colitis[12] and experimental chemotherapy-induced pancreatitis[13]; with *gcn2-*deficient mice showing increased weight loss, higher levels of inflammatory cytokines and worse histological changes compared to their wild-type counterparts. This is intriguingly analogous to our observations – that *gcn2^-/-^* mice showed greater weight loss, higher levels of IL-6 and KC, more severe histological pulmonary changes and increased pulmonary artery pressures after exposure to LPS, which is a classic instigator of inflammation. It is also interesting to speculate that this may be due to the common embryological origins of both lung and gut from the primitive gut tube[37]. For the Ravindran[12] and Keil[14] papers the same transgenic *gcn2^-/-^*mouse line was used as in this manuscript (B6.129S6-Eif2ak4tm1.2Dron/J[38]). However, this susceptibility to inflammation is not peculiar to just this line but is common to *gcn2-*deficiency as a whole, as mice with a missense mutation in *gcn2* (*atchoum,* annotated as *atc,* which results in a near-complete loss of the protein) have defective responses to mouse cytomegalovirus infection, to the point of significant lethality in *gcn2^atc/atc^*mice where none is observed in the wild-type.

We note that the published evidence is mixed when it comes to the effect of GCN2 deficiency in other models of heart failure. *Gcn2* deficiency was protective in a pressure overload model due to transverse aortic banding, with *gcn2-*mutant mice showing lower levels of markers of autophagy and oxidative stress, less left ventricular fibrosis and greater preservation of ejection fraction[39]. Similarly, *gcn2-*deficient mice were protected from hypoxia-induced pulmonary vasoconstriction [40], with Zhu *et al* suggesting that endothelin-1, a potent vasoconstrictor, is a downstream target of GCN2 signalling from both RNA sequencing of wild-type *versus gcn2-*deficient hypoxic mouse lung tissue and from human lung microvascular endothelial cells.

Taken as a whole this leads us to conclude that *gcn2*-deficiency predisposes to inflammation in multiple organ systems, which can lead to specific patterns of end-organ damage. It is fascinating that the only significant phenotype of *gcn2* mutations in patients is pulmonary veno-occlusive disease. This may tie in with our single-cell RNA sequencing results, which indicate that adventitial fibroblasts are likely to be a cell type of interest in GCN2-mutation dependent pulmonary vascular disease. Adventitial fibroblasts have recently been found to be critical drivers of vascular remodelling both in general [41] and with regards to pulmonary arterial hypertension [42, 43]. They are also intimately involved in vascular wall inflammation [44, 45] and so may form the link between *gcn2*-deficiency, inflammation and vascular remodelling.

Examining the adventitial fibroblast cluster in greater detail has also revealed intriguing candidate genes for further interrogation. *Adamts16* is upregulated in *gcn2^-/-^* adventitial fibroblasts compared to the wild-type. It encodes a metallopeptidase which has been linked to blood pressure control in both human genetic studies[46, 47] and in rat models[48]. *Adamts16* activation has also been shown to promote cardiac hypertrophy, fibrosis and subsequent heart failure in a transverse aortic constriction model by activating TGFβ signalling[49]. Similarly, overexpression of *Adamts16* in a rat model of myocardial infarction (due to left anterior descending artery ligation) caused a more severe phenotype with decreased left ventricular systolic pressures and greater apoptotic indices in myocardium[50].

This work opens up potential ways of treating *gcn2-*mutation associated pulmonary vascular disease; and also provides potential insight into how IL-6 may operate in the failing heart. We hypothesize that either blocking IL-6 or its associated pathways may be helpful; or augmenting the integrated stress response (which can be activated via GCN2) using small molecules. The first approach has the advantage of being straightforward but runs the risk of increasing the likelihood of infections[51]. There is a growing body of evidence that IL-6 is implicated in both atherosclerotic cardiovascular disease[52] and heart failure with preserved ejection fraction[30, 31, 53, 54]. Indeed, patients receiving tocilizumab (a recombinant humanised monoclonal antibody that binds to the IL-6 receptor) for treatment of their rheumatoid arthritis showed an increase in their left ventricular ejection fraction and a fall in the left ventricular index assessed by cardiac magnetic resonance imaging[55]. IL-6 inhibition using ziltivekimab (a fully human monoclonal antibody that binds to interleukin-6) reduced biomarkers of inflammation and thrombosis (CRP, fibrinogen and serum amyloid A) in adult patients with chronic kidney disease and high atherosclerotic risk[56, 57]. A randomised control trial is underway to see if ziltivekimab alleviates symptoms and improves function in patients with heart failure (https://clinicaltrials.gov/study/NCT06200207).

The second approach, which aims to restore balance by modulating the integrated stress response, is intriguing as it may avoid the immunosuppressive effects of anti-IL-6 approaches, and would likely temper other mechanisms through which *gcn2* deficiency affects the pulmonary vascular tree and myocardium, thus making its potential applications wider. Both Zhu *et al* [40] and ourselves have shown that *Edn1,* encoding endothelin-1, a potent vasoconstrictor, is affected by the presence of *gcn2.* Pu *et al*[58] have shown that *gcn2* knockdown affects markers of oxidative stress and inflammation in a hypoxia-reperfusion model using rat myocardial myoblasts. Prabhakar *et al*[59] used a rat model of MMC to demonstrate that the resultant pulmonary vascular disease was dependent on activation of protein kinase R and the integrated stress response (ISR), which disrupted the endothelial barrier. From these studies we can surmise that it is highly likely that *gcn2* and the ISR exerts multiple effects through inflammatory, vasoactive, oxidative and metabolic pathways in the pulmonary vascular tree and the myocardium; and that understanding these pathways and their interactions would be a key step in modulating the ISR as a treatment for heart failure.

To summarise we have shown for the first time that *gcn2*-deficiency in mice predisposes to an exaggerated pro-inflammatory response to both lipopolysaccharide and mitomycin C, with the end result of pulmonary vascular remodelling and raised pulmonary vascular pressures. Genetic ablation of *Il6* abrogates this response in both murine models. Both models are accessible and directly reflect patient disease (either genetic loss of GCN2 or exposure to mitomycin C). Single-cell RNA sequencing suggests that adventitial fibroblasts may be a key cell type in this process. Taken together we hypothesize that IL-6 dependent pathways may be an attractive pharmacological target for PVOD, especially as no medical treatments exist for this deadly disease.

## Methods

### Mouse models of pulmonary hypertension

All mouse experiments were formally powered using estimates of variance and minimum detectable differences based on previous experience in genetically modified mouse models of pulmonary hypertension[20]. All animal studies were conducted in accordance with the UK Animals (Scientific Procedures) Act 1986 under the Home Office project licences 70/8550 (NWM), 7550697 (NWM) and 5789170 (SJM).

#### Transgenic mice

*Gcn2^-/-^* mice (B6.129S6-Eif2ak4tm1.2Dron/J, https://www.jax.org/strain/008240) are homozygous for a deletion of exon XII (606-648), which removes an essential portion of the kinase domain. They were obtained from the Jackson Laboratory (Maine, USA), after being initially bred by Prof. David Ron[38]. *Il6^-/-^* mice (B6.129S2-Il6tm1Kopf/J) were the kind gift of Prof. Mendez-Ferrer and Dr Eva Felez, and had previously been obtained from the Jackson Laboratory (https://www.jax.org/strain/002650) and originally developed by Kopf *et al*[60]. The double-deficient mouse was bred in-house by crossing *gcn2^-/-^* and *Il6^-/-^* mice. The genotypes of mice were determined using the automated genotyping service offered by Transnetyx (Tennessee, USA).

#### Baseline phenotyping

Aged (4.5-6 months) mice were anaesthetised using inhaled isoflurane (2.0-2.5%) and subjected to either echocardiography, right and/or left heart catheterisation. For the latter two, pressures and volumes were recorded using the Millar SPR-139 mouse catheter (Millar Instruments, Texas, USA) and subsequently analysed using LabChart (ADInstruments, Colorado, USA). After these procedures, mice were exsanguinated and blood and tissue collected. The right ventricle was dissected and weighed, as was the remaining left ventricle and septum. The right lung was snap-frozen in liquid nitrogen for RNA extraction and the left lung inflated with 4% paraformaldehyde before embedding into paraffin blocks.

#### Acute and chronic mitomycin C models

In the acute experiment, 8–10-week-old wild-type mice were exposed to either intraperitoneal vehicle or 1.0mg/kg of mitomycin-C from *Streptomyces caespitosus* (M7949, Sigma-Aldrich, UK) and humanely killed at 3- and 24-hours post-exposure for collection of blood and tissue as described above. In the chronic experiment, 8–12-week-old wild-type, and *il6^-/-^* mice were injected with either 0.5mg/kg, 1.0mg/kg, or 2.0mg/kg (in total) of mitomycin-C or vehicle and then subjected to right heart catheterisation and tissue collection under terminal anaesthesia as described above (Fig.4A).

#### Acute and chronic LPS models

In the acute experiment, 8–10-week-old wild-type mice were exposed to either intraperitoneal vehicle or 0.05mg/kg or 0.1mg/kg of LPS (*E.coli* 0111:B4, Sigma-Aldrich, UK) and humanely killed at 3 hours post-exposure for collection of blood and tissue as described above. For the chronic experiment, 8-12 week old wild-type, *gcn2^+/-^*, *gcn2^-/-^*, *Il6^-/-^* and *gcn2^-/-^Il6^-/-^* mice were exposed to either intraperitoneal vehicle or 0.05mg/kg or 0.1mg/kg of LPS (*E.coli* 0111:B4, Sigma-Aldrich, UK) thrice weekly for six weeks and then subjected to right heart catheterisation and tissue collection under terminal anaesthesia as described above.

#### Assessment of pulmonary vascular muscularisation in mouse lung

Paraffin embedded mouse lung tissue sections were baked at 60°C for an hour, dewaxed and rehydrated, on the ST5020 Workstation (Leica, Germany). The staining for alpha-smooth muscle actin was performed on a Leica automated Bond-III platform in conjunction with the Polymer Refine Detection System (DS9800) using the Abcam antibody (catalogue number ab5694, Abcam, UK) and the DAB enhancer (AR9432, Leica, Germany). This work was done by the Histopathology and In-Situ Hybridisation core facility at the Cancer Research UK Cambridge Institute.

Pulmonary vessel muscularisation was quantified by identifying small vessels (50-200 microns in diameter) in a 10mm^2^ lung section per mouse and classified by a one of three blinded observers as to whether they were non-muscularised, partially muscularised or fully muscularised, depending on the degree of smooth muscle actin staining. Examples of this grading can be found in Supplementary Methods. Statistical significance was determined using Fisher’s contingency testing to compare the percentage of non-muscularised: percentage of partially muscularised: percentage of fully muscularised vessels between groups.

#### Quantitative PCR

RNA was extracted from tissue using TRIzol (ThermoFisher Scientific, Massachusetts, USA) and then subjected to purification with the RNeasy Mini kit with DNAse digestion (Qiagen, Venlo, the Netherlands). The RNA was reverse-transcribed with an Applied Biosystems cDNA reverse transcription kit (Thermo Fisher Scientific, Waltham, MA) to generate cDNA. Q-PCRs were prepared with SYBR green JumpStart Taq ReadyMix (Sigma-Aldrich). Reactions were amplified on a C1000 Touch Thermal Cycler (BioRad, Watford, UK) using primers from Qiagen and Sigma-Aldrich. The expression of target messenger RNA (mRNA) was normalised to beta-actin, using the DDCT method. Further details are available in the online supplement.

### Cell models and treatments

Mouse embryonic fibroblasts were generated in-house from *gcn2^-/-^*and their wild-type littermates. After treatment with lipopolysaccharide (*E. coli* 0111:B4, Sigma-Aldrich, UK) or mitomycin C from *Streptomyces caespitosus* (M7949, Sigma-Aldrich, UK) or vehicle (media alone), they were lysed, RNA extracted using RNeasy Mini kit, and quantitative PCR conducted as above. Alternatively, the conditioned media was collected and used for quantification of secreted IL-6 as below.

### Quantification of IL-6 levels in supernatants

Mouse IL-6 levels in conditioned media from MEFS were assayed using an in-house ELISA. Further details are available in the online supplement.

### Patient studies

#### Patient identification

Newly diagnosed patients with a confirmed diagnosis of pulmonary veno-occlusive disease were prospectively recruited at the national pulmonary hypertension centres in the UK from 2014 onwards as part of the National Cohort Study of Idiopathic and Heritable PAH (ClinicalTrials.gov identifier: NCT01907295). The study was approved by the local research ethics committee (references 13/EE/0325 and 13/EE/0203) and all participants gave written informed consent. Patients who had been previously diagnosed were also approached and asked for consent to be included in the cohort. Pulmonary veno-occlusive disease was defined as having a mean PA pressure of >25mmHg with a capillary wedge pressure of <15mmHg and having either homozygous or compound heterozygous mutations in *EIF2AK4*.

#### Plasma sampling and cytokine measurements

Blood sampling was performed at the time of the first patient visit. Venous blood was collected in plasma-citrate tubes, inverted 3-4 times, and centrifuged within 40-60 minutes at 1500x g for 15 minutes. The samples were then aliquoted and stored at -80°C until analysed. Patient IL-6 levels were measured using a custom assay manufactured by Meso Scale Discovery (Maryland, USA). This was performed by staff in the Core Biochemical Assay Laboratory at Cambridge University Hospital, UK, who were blinded to the identity and genotypes of the samples. Further details are available in the online supplement.

### Single-cell RNA sequencing of mouse lungs

#### Protocol

Age- and sex-matched *gcn2^-/-^* and wild-type littermates from the same cage were sacrificed by cervical dislocation. The lungs were flushed with PBS-heparin (25U/ml), washed thrice in PBS, and finely minced. The lung homogenates were digested (using collagenase, dispase and DNase), filtered through a 100micron sieve, subjected to red blood cell lysis, and then passed through a 50micron filter. Flow cytometry was used to isolate single live CD45-positive and CD45-negative cells and 16,000 cells were processed using the Chromium 3’ kit (10xGenomics, Leiden, the Netherlands). Sample processing and cDNA library preparation was performed by the Jeffrey Cheah Biomedical Centre and the Cancer Research UK genomics core facility. The cDNA was sequenced on a NovaSeq S2 or S4 chip (NovaSeq 6000, Illumina, UK). Further details are available in the online supplement.

#### Bioinformatics

This was performed in collaboration with Dr Kishore at the Cancer Research UK Cambridge Institute. Briefly, the raw sequences were aligned with Cell Ranger using the STAR aligner to the reference mouse genome -GRCm38.p6[61]. Filtering and quality control assessments were performed using the SCATER R package and dimensionality reduction was performed using the PCA, T-SNE and UMAP algorithms[62, 63]. Cells were clustered into similar types using the Walktrap community detection algorithm, and top marker genes for each cluster were then identified[64, 65]. A gene-set variation analysis was performed with these markers to identify the cell type clusters, using previously-identified cell markers[66]. A ‘pseudo-bulk’ method to aggregate cells together from the same biological replicate was then used to identify differentially expressed genes between wild-type and *gcn2^-/-^* cells in each cluster[25, 67].

### Statistics

Statistical analysis was performed using GraphPad Prism version 10 (USA), R and Python. Datasets were tested for adherence to a normal distribution using the Shapiro-Wilks test. Data was compared using unpaired t-tests (for normal data), Mann-Whitney tests (for non-normal data) and Fisher’s (for categorical data). For mouse experiments, power calculations were performed using an online power calculator made available by the University of Illinois (USA) and the numbers of mice needed for a power of 90% and a significance level of <0.05 were calculated. The Home Office regulations state that the number of experimental mice used must be kept to a minimum consistent with study aims (Animal [Scientific Procedures] Act 1986, https://www.gov.uk/government/collections/animals-in-science-regulation-unit). Therefore, a pre-planned analyses was performed at the sample size needed for a power of 80%. If the result was robust, then further numbers were deemed unnecessary.

## Data sharing

The raw data for the single cell RNA sequencing is being submitted to the Gene Expression Omnibus database (https://www.ncbi.nlm.nih.gov/geo/). The accession number will be available for reviewers if needed in 5-7 working days. All other data is available on reasonable request to the corresponding author.

## Acknowledgements

We acknowledge the help and support of Professor David Ron (University of Cambridge); Ms Maike Paramor, Ms Kania Katarzyna and members of the CRUK and JCBC NGS facilities; and members of the Marciniak and Morrell laboratories. Special thanks to all the patients with PVOD, healthy volunteers, and staff in the pulmonary vascular centres in the UK who have selflessly given their tissue, data, and time towards this effort.

## Author contributions

MSch designed and performed experiments, analysed and interpreted data and helped write the manuscript. QG, MSou, AC, SM, JAV, BD, and NV performed experiments and analysed and interpreted data. PU, RT, NWM and SJM provided critical laboratory inputs, helped plan experiments, interpreted data, and helped write the manuscript. All authors were involved in reading and critically revising the manuscript. ES designed and performed experiments, analysed and interpreted data, wrote the manuscript, and funded the study.

