## Supplement for "Interleukin-6 is critical in the development of *gcn2-*mutation associated pulmonary vascular disease in mice"

**Supplementary Methods**

**Quantitative PCR**

Mouse embryonic fibroblasts (MEFs) derived from *gcn2^-/-^* or wild-type littermate embryos were plated in 6-well plates for RNA studies at densities of 3 x10^5^ cells/well. After treatment, cells were lysed and RNA extracted using the RNeasy Mini Kit (Qiagen). The subsequent steps were performed in exactly the same way as tissue RNA. The primers used are shown below in Supplementary Table 1.

| **Gene** | **Species** | **Forward 5’-3’** | **Reverse 5’-3’** | **Source** |
| --- | --- | --- | --- | --- |
| *Actin* | Mus musculus | TCCTGGCCTCACTGTCCA | GTCCGCCTAGAAGCACTTGC | W’out *et al* |
| *Chop* | Mus musculus | GGAGCTGGAAGCCTGGTATGAG | GCAGGGTCAAGAGTAGTGAAGG | W’out *et al* |
| *Il6* | Mus musculus | Qiagen-proprietary  Mm_IL6_1_SG | Qiagen-proprietary |  |
| *Cxcl1* | Mus musculus | Qiagen-proprietary  Mm_Cxcl1_1_SG | Qiagen-proprietary |  |
| *Eif2ak4* | Mus musculus | Qiagen-proprietary  Mm_Eif2AK4_1_SG | Qiagen-proprietary |  |

**Supplementary Table 1: Primers used for quantitative PCR**

**ELISA**

For cell experiments, mouse embryonic fibroblasts (MEFs) were plated in 24-well plates for ELISA studies at densities of 3 x10^4^ cells/well. After exposure to either the treatment or to vehicle only the supernatants were harvested and the cells in the wells counted for normalization purposes. Flat-bottomed medium-binding 96-well plates (Nunc-Immuno, M9140, Sigma-Aldrich, UK) were coated with 50 microlitre of capture antibody (Supplementary Table 2) diluted in a carbonate buffer for 2 hours at room temperature. The plate was then washed thrice using PBS containing 0.05% Tween (v/v, PBS-T) and blocked using 5% foetal bovine serum in PBS-T for an hour. 50microlitres of samples and standards were added and the plate incubated overnight at 4oC. After three washes with PBS-T, a biotinylated capture antibody was added and left to incubate for 2 hours. The plate was then washed three times as before and incubated with ExtraAvidin alkaline phosphatase conjugate (Sigma-Aldrich, UK) at 1/400 for a further 2 hours. After a further 2 washes with PBS-T and a final wash with distilled water, the substrate P-nitrophenylphosphate (PNPP, Sigma-Aldrich, UK) was added at a concentration of 1 microgram/ml in diethanolamine buffer (10mM diethanolamine, 0.5mM MgCl2). The absorption at 405nm was measured using a Tecan CM Spark plate reader (Tecan, Switzerland). A four-parameter logistic curve was fitted to the standards and used to interpolate the concentration of the unknown samples. Protein levels were then normalised to the number of live cells in the corresponding well.

| **Antibody** | **Species recognised** | **Manufacturer** | **Concentration of antibody** | **Vehicle** |
| --- | --- | --- | --- | --- |
| IL-6 | Mouse | R&D Systems  MAB406 | 2ug/ml | Carbonate coating buffer |
| IL-6 biotinylated | Mouse | R&D Systems  BAF406 | 0.3µg/ml | 5% foetal bovine serum in PBS-Tween |

**Supplementary Table 2: Antibodies used for ELISA**

**Immunohistochemistry**

After the lung slides had been stained, they were scanned using a Hamamatsu NanoZoomer XR and analysed using NDP.view2 software (<https://www.hamamatsu.com/eu/en/product/life-science-and-medical-systems/digital-slide-scanner/U12388-01.html>, Hamamatsu Photonics, Japan). They were then classified by one of three blinded reviewers as being non-muscularised, partially muscularised or fully muscularised, dependent on the degree of smooth muscle-actin staining (Supplementary Table 3).

| **Category** | **Degree of muscularisation** | **Representative example** |
| --- | --- | --- |
| Non-muscularised | ≤10% smooth-muscle actin staining | 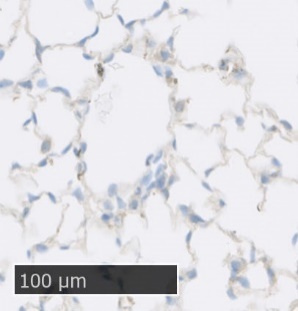 |
| Partially muscularised | >10-90% smooth-muscle actin staining | 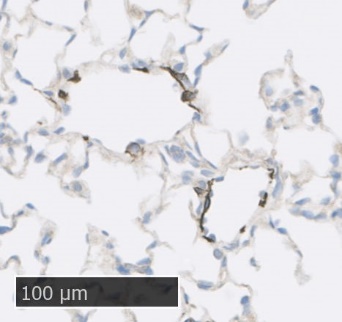 |
| Fully muscularised | >90% smooth-muscle actin staining | 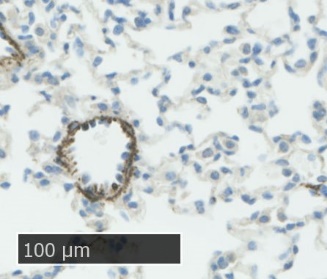 |

**Supplementary Table 3: Definitions used for pulmonary vessel grading**

**Patient genotyping**

All patients had next-generation paired-end whole genome sequencing using Illumina HiSeq2500 and HiSeqX (Illumina, San Diego, USA). DNA was extracted from whole blood at the central extraction and QC laboratory in Cambridge. Reads were aligned against the Genome Reference Consortium human genome build 37 (GRCh37, GenBank:2648) using the Illumina Isaac Aligner (version SAAC00776.15.01.27) and variants were then called using the Illumina Starling Software (version 2.1.4.2)[1]. The variants were then left-aligned, normalised with BCFtools and loaded into our Hbase database (Wilmington, USA) to produce multi-sample variant calls for the genetic association studies[2].

**Cytokine measurements in patient plasma**

A custom multiplex assay manufactured by Meso Scale Discovery (MSD, Maryland, USA) was used to measure cytokine and growth factor levels in patient plasma. Calibration curves were prepared in the assay diluent, with a range of 1.2 to 40,000pg/ml, dependent on the cytokine. Arrays were pre-incubated with 25 microlitres per well of assay diluent for 30 minutes. 25 microlitres of sample or calibrator were added in duplicate to wells in the plate and then incubated at room temperature for 2 hours. The array was washed with PBS-T, and 25 microlitres of detection antibody was added. After a further 2 hours of incubation at room temperature, the array was washed and the detection solution added. Results were read using an MSD Sector Imager 6000 (Maryland, USA). Cytokine concentrations were determined with SoftMax Pro (version 4.6, Molecular Devices, California, USA) using curve fit models.

**Single cell RNA Sequencing**

A pilot experiment was initially conducted to optimise the protocol for harvesting mouse lungs and dissociating them into suspensions containing only live single cells representative of the lungs from which they were sourced. This revealed that >90% of cells isolated were CD45-positive. Therefore an additional step was introduced to increase the proportion of non-immune cells so they could be characterised, particularly as endothelial cells and smooth muscle cells are usually thought to be the main cell types of interest in pulmonary hypertension[3, 4]. The single cell suspension was subjected to flow cytometry analysis and sorted based on the haematopoietic marker CD45 (Alexa Fluor® 488 anti-mouse CD45 Antibody, BioLegend, UK) to separate immune and non-immune cells and ensure that the same numbers of each were submitted to allow for single cell RNA sequencing of both compartments (Supplementary Figure 1).

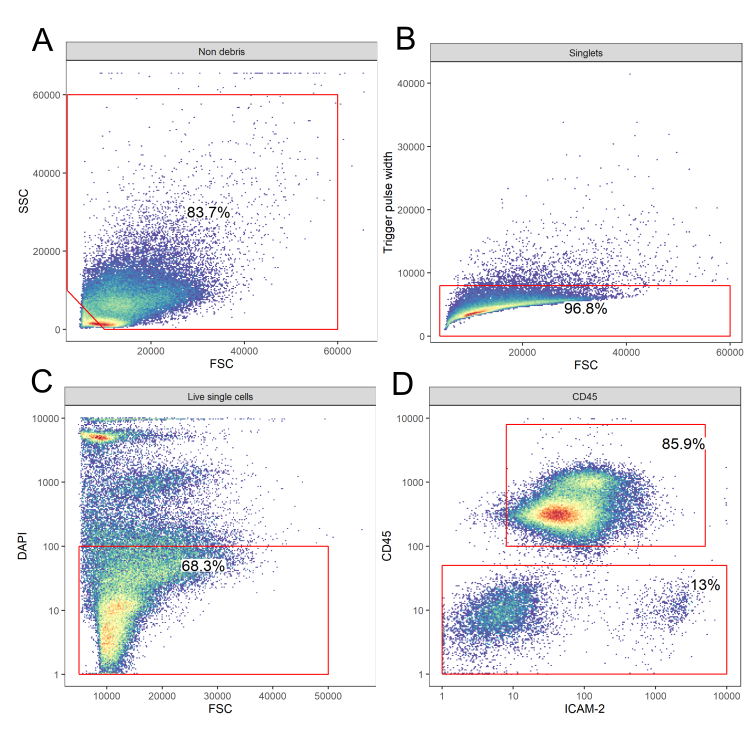

**Supplementary Methods Fig. 1: Flow cytometry analysis and sorting for single cell RNA sequencing samples.**

Panels A-C show gating used to eliminate debris (panel A), to isolate single cells (panel B), and to sort for live (*i.e.* DAPI-negative) cells (panel C). Panel D shows gating for CD45-positive (blue) and CD45-negative (pink) cells, with additional ICAM-1 staining to visualise the endothelial cell population.

**Supplementary Results**

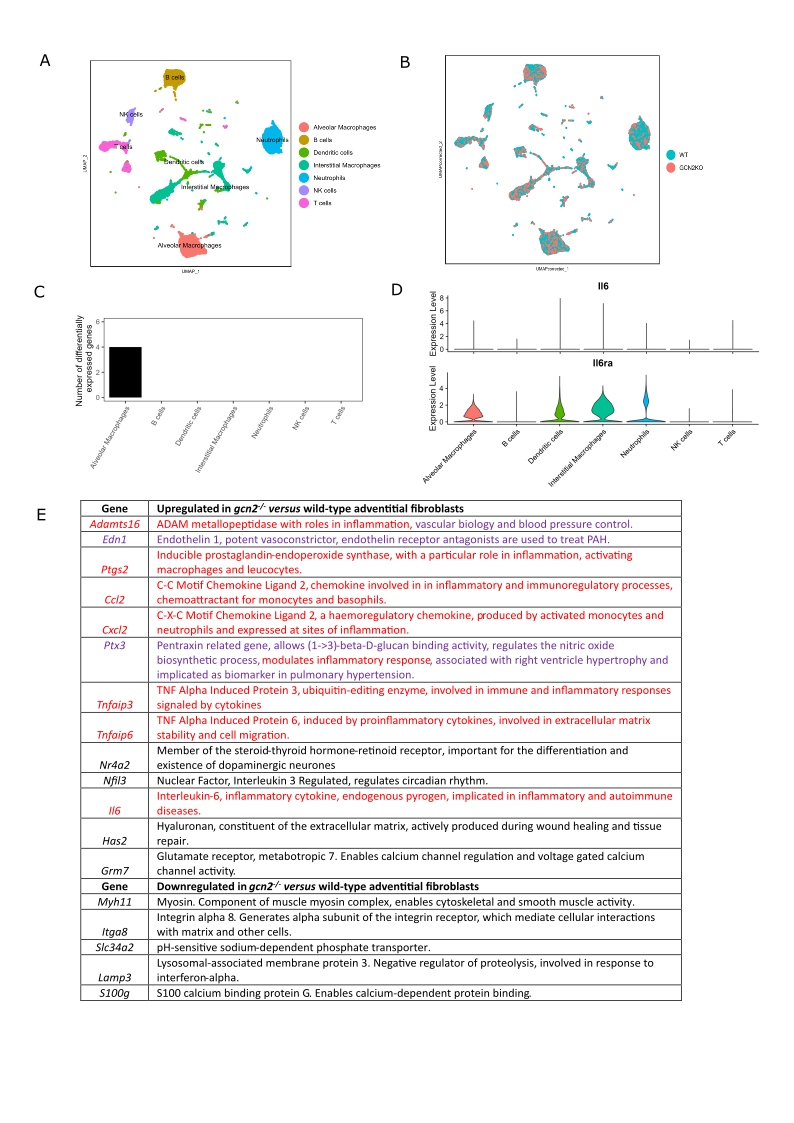

**Supplementary Figure 1: Further elucidation of single-cell RNA sequencing datasets**

Panels A and B show the Uniform Manifold Approximation and Projection graphs (UMAPs) of CD45-positive lung cells, with panel A depicting cell clusters and panel B showing contributions from the wild-type and *gcn2^-/-^* mice. Panel C shows the cell types from the CD45-negative group arranged in order by the number of genes differentially expressed between wild-type and *gcn2^-/-^* lung cells. Panel D shows violin plots showing the relative expression of *Il6* and *Il6ra* in cell types from the CD45-positive dataset. Panel E shows the genes most upregulated and downregulated in *gcn2^-/-^* versus wild-type adventitial fibroblasts and a summary of their roles. Genes encoding proteins which have inflammatory roles are in red while genes encoding proteins with known roles in pulmonary hypertension, systemic hypertension or vascular biology are in purple.

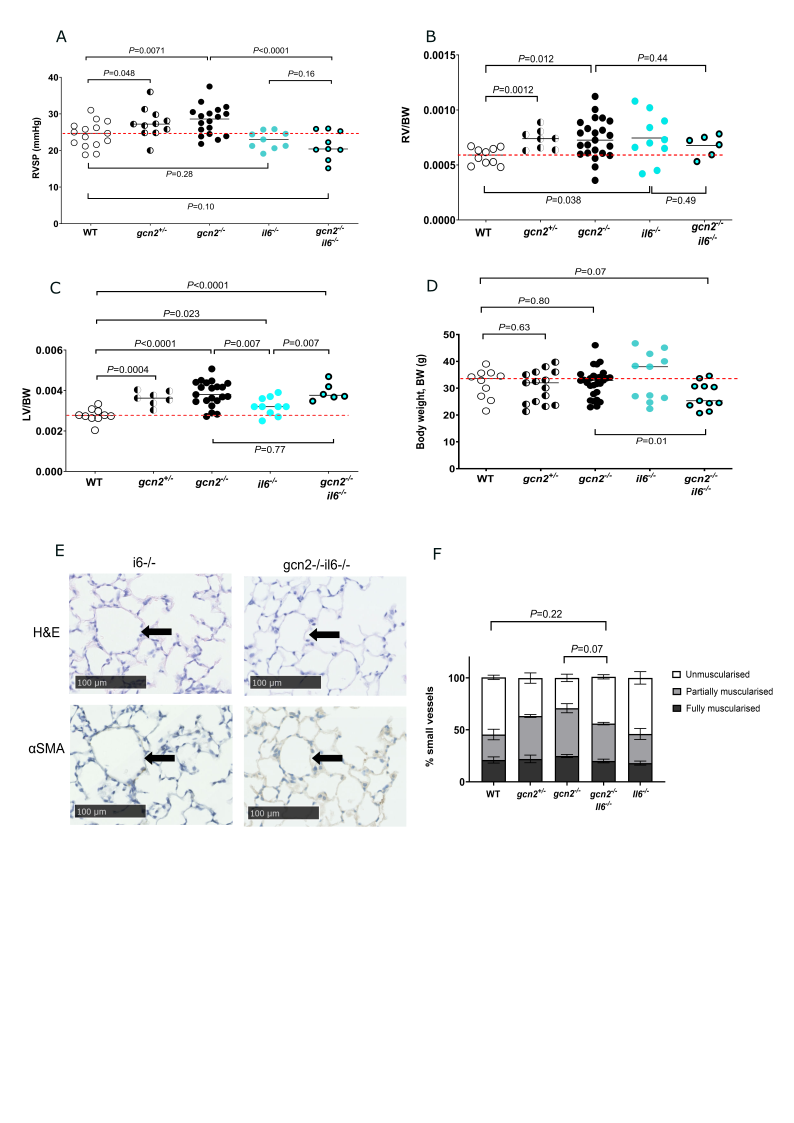

**Supplementary Figure 2: Characterisation of the *il6*-deficient mouse and the *gcn2^-/-^ il6^-/-^* mouse at baseline.**

Panels A-D show the right ventricular systolic pressure (RVSP, panel A), the right ventricular mass indexed to body weight (RV/BW, panel B), the left ventricular mass indexed to body weight (LV+S/BW, panel C) and the body weight (panel D) in il*6^-/-^* and *gcn2^-/-^ il6^-/-^* mice. Note that this panel includes mice whose data has been previously shown in Figure 1 (for wild-type, *gcn2^+/-^* and *gcn2^-/-^* mice) as by UK law we are required to use the minimum number of mice to fulfil the needs of an experiment. Panels E-F show representative histological sections of the lungs of il*6^-/-^* and *gcn2^-/-^il6^-/-^* mice, stained with haematoxylin and eosin (H&E) and smooth muscle actin (SMA) and the quantification of non-muscularised, partially muscularised and fully muscularised vessels (panel F). In panels (A-D) the median is shown, and comparisons have been made using unpaired t-tests (for parametric data) or Mann-Whitney tests (for non-parametric data). In panel F the distribution of non-muscularised: partially muscularised: fully muscularised vessels has been compared between groups using Fisher’s contingency testing.
